# The response of the skeletal muscle transcriptome to exercise in chronic kidney disease

**DOI:** 10.1101/2025.04.25.650585

**Authors:** Luke A Baker, Matthew Graham-Brown, Thomas J Wilkinson, Alice C Smith, Emma L Watson

## Abstract

**Background:** Chronic kidney disease (CKD) affects approximately 14% of the UK population and is associated with significant exercise intolerance, partly due to skeletal muscle dysfunction. While exercise is a potential therapeutic strategy, the molecular response of skeletal muscle to exercise in CKD remains poorly understood. This study aimed to characterise transcriptomic changes in skeletal muscle 24 hours after aerobic (AE) or combined aerobic and resistance exercise (CE) in non-dialysis CKD.

**Methods:** This study utilised muscle biopsies from participants in the ExTRA CKD trial with stage 3b–4 CKDages 3b-4 (AE: 24 (15-32) ml/min/1.73m^2^; CE: 25 (19-31) ml/min/1.73m^2^). Participants (n=4 per group) were randomised to 12 weeks of thrice-weekly AE or CE. Vastus lateralis skeletal muscle biopsies were collected at baseline and 24h after the first bout of exercise. RNA was extracted for Bulk RNA sequencing. Bulk RNA sequencing was performed, and differentially expressed genes (DEGs) were identified between baseline and post-exercise samples, followed by pathway enrichment analysis.

**Results:** Following AE, 1480 genes were upregulated and 1554 downregulated. CE resulted in 556 upregulated and 115 downregulated genes. The most upregulated gene after AE was *CHI3L1* (log₂FC 10.7), followed by *SAA2* and *PTX3*, all associated with inflammation. After CE, *SFN* (log₂FC 6.8) and *MT1A* were among the most highly upregulated. Enrichment analysis showed strong activation of inflammatory and cellular senescence pathways, and downregulation of mitochondrial function-related processes, particularly after AE.

**Conclusion:** Both AE and CE triggered robust inflammatory gene expression responses in CKD skeletal muscle, indicative of early repair processes. Unexpectedly, mitochondrial-related pathways were downregulated, aligning with previous findings of impaired mitochondrial adaptation in CKD. These results highlight mitochondrial dysfunction as a potential barrier to effective exercise adaptation and a possible therapeutic target in this population.

## Introduction

Chronic kidney disease (CKD) is a growing public health emergency, affecting around 14% of adults in England and is projected to increase in the coming decades (1). CKD patients have a high symptom burden that includes exercise intolerance, resulting in high levels of physical inactivity (2). There are several possible reasons for this poor exercise tolerance. One contributing factor to this is peripheral limitations, for example, skeletal muscle wasting and weakness, which are also common in people with CKD (3).

It is now well established that in people with CKD skeletal muscle dysfunction is common, which includes muscle atrophy, muscle weakness (4), and maladaptive molecular changes including insulin resistance (5) and intramuscular inflammation (6). Previous studies have shown that skeletal muscle also elicits an abnormal response to exercise, which reports a blunted anabolic (7) and mitochondrial biogenesis (8) in response to exercise, together with a slower recovery of phosphocreatine which suggests reduced mitochondrial function (9). However, data in this area is scant and we do not fully understand the molecular events that occur within skeletal muscle following acute exercise or exercise training in the CKD population. This understanding is vital to guide the optimisation of strategies to improve physical function, overall health and quality of life and support adaptations to exercise. Technological advances now enable a myriad of exploratory analysis methodologies which allow for the unbiased investigation of the human transcriptome (10). To date, no studies have used an untargeted approach to investigate changes to the skeletal muscle transcriptome in response to exercise in chronic kidney disease. Therefore, this study aimed to report key changes in the skeletal muscle transcriptome 24h after aerobic exercise (AE) or combined aerobic and resistance exercise (CE) in non-dialysis CKD.

## Methods

### Study design and participants

Participants included in this study (ExTra-CKD trial (ISRCTN: 36489137)) had CKD not yet requiring dialysis and had consented to donate skeletal muscle biopsies. The main trial paper has been published previously (11). The recruitment period started 16^th^ December 2013 to 30^th^ April 2016 with the intervention period completed in October 2016. Exclusion criteria included: age <18 years, physical impairment sufficient to prevent undertaking the intervention, recent myocardial infarction, unstable chronic conditions, or an inability to give informed consent, and a BMI >40 (due to difficulties in muscle size measurement). Diabetic patients were included if haemoglobin A1C was <9%. In summary, participants were randomised using a random block method stratified for CKD stage, to either 12-weeks 3x/week supervised CE, or AE alone to a thrice-weekly 12-week programme of either aerobic exercise only (AE) only or combined exercise (aerobic + resistance exercise; CE). The randomisation sequence was generated by using online software. The AE component consisted of circuits of up to 30 minutes of treadmill, cycling and rowing exercises at 70-80% heart rate maximum. The CE group performed the same aerobic exercise component but with the addition of 3 sets of 12-15 repetitions of leg extension and leg press exercises on a fixed weights machine at 70% 1-repetition maximum. The amount of work performed between the two groups was matched for duration. Ethical approval was granted from the UK National Research Ethics Committee (13/EM/0344). All participants gave written informed consent, and the trial was conducted per the Declaration of Helsinki.

### Sample collection, processing and RNA sequencing

Muscle biopsies were collected from the vastus lateralis after an overnight fast using the needle biopsy technique at two timepoints: baseline and 24h after the first exercise session. Following the dissection of any visible fat, samples were immediately placed in liquid nitrogen and stored until subsequent analysis.

RNA was isolated from approximately 10mg/wet weight muscle biopsy tissue using TRIzol® (Fisher Scientific, UK) per the manufacturer’s instructions. RNA quality was assessed using the Ailgent 5400 fragment analyser system. The mean RIN value for RNA submitted for sequencing was 8.0+/− 2.1. All library preparation and subsequent RNA sequencing was performed by Novogene (Beijing, China) using the Illumina Novoseq 6000 sequencing system using a 150bp paired end strategy. HISAT2 was used to align the sequencing reads to the reference genome (Homo sapiens GRCG38/hg38).

### Statistical analysis

The abundance of each gene was quantified using feature counts and then normalised to the total number of reads for each gene. The data was filtered and reads with adapter contamination, reads when it was uncertain nucleotides constitute more than 10% of either read, and reads with low quality nucleotides that constitute of more than 10% of either read were removed. The reads were aligned to a reference genome (Homo Sapiens(GRCh38/hg38)) using HIAT2. To determine reliability of the analysis, correlation of the gene expression levels between samples was determined by Perasons correlation. Principal component analysis (PCA) was used to evaluate intergroup differences and intragroup sample duplication. Due to the presence of biological replicates differential gene expression (DEG) analysis was performed using the DESeq2 package in R using the negative binomial distribution model. Hierarchical clustering analysis (HCA) was conducted to cluster differentially expressed genes (DEGs) and visualize their expression profiles pre and post exercise in the different groups. Sequencing depth and RNA quality were added into the analysis as covariates to control for potential confounding effects. Genes with adjusted P value <0.05 and with a |log2(FoldChange)|>=1 and a false discovery rate (FDR) of 0.05 (12) (using the Benjamini-Hochberg procedure) were considered differentially expressed. A hierarchical clustering analysis of DEGs was performed using the ggplot2 package in R. Functional enrichment analyses including Gene Ontology (GO) and Kyoto Encylopedia of Genes and Genomes (KEGG) pathway analysis were performed using the clusterProfiler software to determine which DEGs were significantly enriched in which GO terms or metabolic pathways. GO terms and KEGG pathways with padj < 0.05 were deemed significantly enriched.

## Results

### Participant characteristics

Eight participants were included in this analysis, n=4 randomised to AE, and n=4 to CE (Table 1). Both groups were well matched for eGFR (AE: 24[15-32]ml/min/1.72m^2^; CE 25[19-31]ml/min/1.72m^2^) and age (AE: 63 [56-78]years; CE 61[52-80]years). The AE group was made up of 50% males and the CE group 75% males. Co-morbidities included type II diabetes, hypertension, ischaemic heart disease, and valvular heart disease.

**Table 1.** Participant characteristics.

|  | **AE group (n=4)** | **CE group (n=4)** |
| --- | --- | --- |
| **Number of Males (%)** | 2 (50%) | 3 (50%) |
| **Age (y)** | 63 (56-78) | 61 (52-80) |
| **BMI (kg/m<sup>2</sup>)** | 27.5 (24-32) | 32.5 (29-39) |
| **eGFR (mL/min/1.73m<sup>2</sup>)** | 24 (15-32) | 25 (19-31) |
| **Haemoglobin (g/dL)** | 122 (102-161) | 111 (103-131) |
| **Albumin (g/L)** | 44.5 (39-47) | 39 (32-43) |
| **Comorbidities (n)** |  |  |
| **Type II Diabetes** | 0 | 1 |
| **Essential hypertension** | 3 | 2 |
| **Ischaemic Heart Disease** | 0 | 1 |
| **Valvular heart disease** | 1 | 0 |

Data are shown as median and range

### Changes in gene expression following an unaccustomed session of exercise

DEG analysis showed that 24h after the first session of AE 1480 genes were significantly upregulated, 1554 genes were downregulated and 24,142 were unchanged (Figure 1A). 24h following one session of CE, 556 genes were significantly upregulated, 115 were downregulated and 25,435 were unchanged (Padj<=0.05; |log2foldchange|>1) (Figure 1B). HCA (Figure 1C) shows the distinct regulation pattern of genes 24h following both AE and CE. In addition, perasons correlation analysis showed all biological replicates had an R^2^ value >0.8.

**Figure 1.**
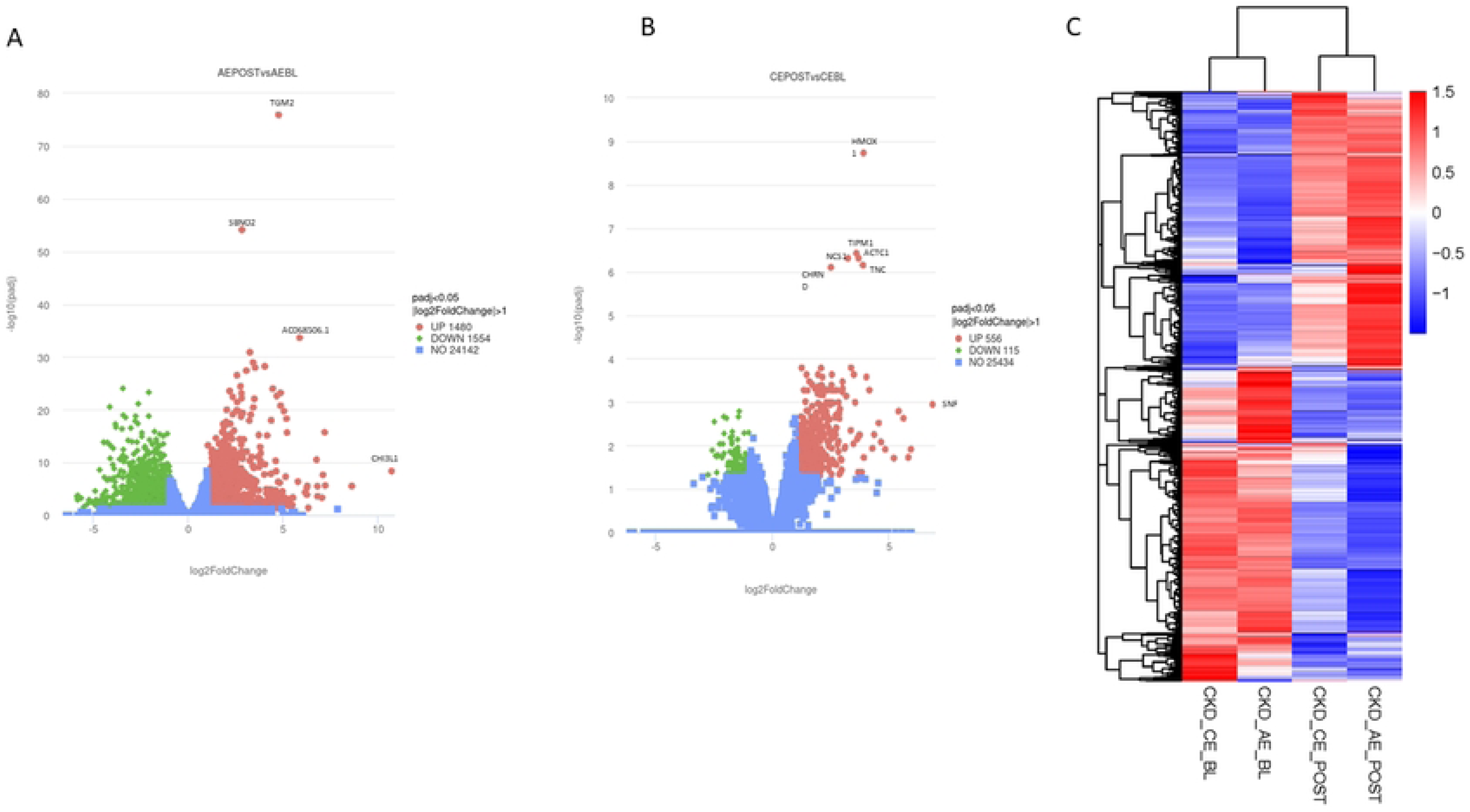
Skeletal muscle transcriptomic response to acute aerobic and combined exercise in CKD. (A) Volcano plot showing differentially expressed genes (DEGs) in vastus lateralis muscle 24 hours after a single session of aerobic exercise (AE) in participants with non-dialysis CKD (n = 4). (B) Volcano plot showing DEGs following combined aerobic and resistance exercise (CE) in a matched CKD cohort (n = 4). In both plots, each point represents a gene, with red indicating significant upregulation, green indicating significant downregulation, and blue indicating no significant change (adjusted *p* < 0.05, |log₂ (fold change)| ≥= 1). Key genes with large fold changes are annotated. (C) Hierarchical clustering analysis (HCA) of DEGs pre- and post-exercise in AE and CE groups, demonstrating distinct gene expression patterns induced by each exercise modality. Heatmap colours indicate scaled expression levels (blue = low, red = high). Clustering confirms differential transcriptomic responses to AE and CE within the CKD muscle environment.

### Skeletal muscle transcriptome response to AE

A list of the top 20 upregulated genes following AE is shown in Table 2. The gene with the largest fold increase in response to AE was CHI3L1 (10.7lfc, P<0.0001), a pro-inflammatory acute phase protein, followed by SAA2 (serum amyloid A2) (8.6lfc, P<0.0001) and PTX3 (pentraxin 3, involved in inflammatory responses) (7.2 lfc, P<0.0001). The most significantly upregulated gene following AE was TGM2 (transglutaminase 2; 4.7lfc, P<0.0001).

**Table 2.** The top 20 upregulated genes following AE.

| Gene Symbol | Gene name | Gene Function | Log2fc | Adj P value |
| --- | --- | --- | --- | --- |
| CHI3L1 | Chitinase 3 like 1 | A glycoprotein that plays a role in inflammatory and immunological disease, including cancer, Alzheimer's disease and atherosclerosis. It is upregulated in response to tissue injury or inflammation potentially influencing the repair and regeneration of muscle fibers by regulating the activity of immune cells like macrophages. | 10.7 | 4.28E-09 |
| SAA2 | Serum amyloid A2 | An acute-phase protein that is produced in response to tissue injury and inflammation. It is a biomarker of inflammation and is involved in cholesterol homeostasis and HDL metabolism | 8.6 | 2.90E-06 |
| PTX3 | Pentraxin 3 | Plays a role in the innate immune system, inflammation and tissue repair. In muscle, PTX3 is upregulated in response to muscle injury or stress and contributes to the inflammatory response. | 7.2 | 2.35E-06 |
| MT1A | Metallothionein 1A | Has roles in protection against reactive oxygen species., regulating physiological balance and immune homeostasis | 7.1 | 1.82E-16 |
| MYL10 | Myosin light chain 10 | Involved in cardiac conduction, smooth muscle contraction and calcium binding | 8.4 | 0.04 |
| IL1RL1 | Interleukin 1 like receptor 1 | A receptor for IL-33 which promotes the activation of various immune cells, including T-helper 2 (Th2) cells, mast cells, basophils, and eosinophils. In muscle, it helps coordinate the immune response to injury. | 7.0 | 0.0004 |
| SFN | Stratifin | Has a crucial in maintaining cell integrity. Functions include cell growth and development, cell cycle regulation, response to cellular stress, a scaffold for protein interactions. In muscle, it modulates oxidative stress, creating a favourable environment for muscle repair and regeneration. | 6.8 | 0.0002 |
| CCL19 | C-C motif chemokine ligand 19 | A chemokine that is involved in immune cell migration, enabling the efficient immune surveillance, activation, and organization of immune cells within lymphoid tissues | 6.3 | 0.04 |
| MFSD2A | Major facilitator superfamily domain containing 2A | Involved in the transport of essential fatty acids such as omega-3-fatty acids | 6.2 | 0.0001 |
| SAA1 | Serum amyloid A1 | An acute phase protein that is upregulated in skeletal muscle in response to injury and acts as a pro-inflammatory mediator initiating the immune response required for tissue repair and remodelling. | 6.2 | 7.28E-06 |
| GAL | Galanin and GMAP prepropeptide | Peptides derived from preprogalanin. They have roles to play in muscle repair and regeneration, fat metabolism. | 6.1 | 4.86E-05 |
| LIPG | Lipase G endothelial type | Also known as endothelial lipase with functions in lipid metabolism and has anti-inflammatory effects. | 5.7 | 4.35E-07 |
| NIPAL4 | NIPA like domain containing 4 | Involved in ion transport and in magnesium homeostasis | 5.5 | 0.0004 |
| HTR1B | 5-hydroxytryptamine receptor 1B | A neurotransmitter with roles in blood flow regulation and muscle contraction | 5.4 | 0.01 |
| SPOCD1 | SPOC domain containing 1 | Involved in chromatin remodelling, gene expression regulation and DNA repair | 5.4 | 0.0007 |
| C3orf52 | Chromosome 3 open reading frame 52 | Precise biological functions are not yet fully elucidated | 5.3 | 0.0009 |
| FCGR1A | Fc fragment of IgG receptor 1a | A receptor of the FC region of IgG antibodies with a primary role in phagocytosis | 5.3 | 0.0008 |
| SOCS3 | Suppressor of cytokine signalling 3 | Negative regulator of cytokine signalling, inhibiting pro-inflammatory cytokines such as IL-6, IL-10 and IL-4. | 5.1 | 5.07E-19 |
| CELC4G | C-type lectin domain family 4 member G | A pattern recognition receptor expressed primarily on dendritic cells and monocytes, resulting in immune cell activation | 5.1 | 0.005 |
| SELE | Selectin E | A cell adhesion molecule expressed on the surface of endothelial cells in response to inflammatory stimuli. In muscle it is involved in the inflammatory response that occurs following injury or stress. | 5.1 | 0.0004 |

Table represents the log fold change values from baseline to 24h following an unaccustomed session of aerobic exercise (AE) together with a brief description of known function. Abbreviations: fc, fold change

A list of the top 20 downregulated genes is shown in Table S1. The gene with the largest fold decrease was GRIN2A (glutamate ionotropic receptor NMDA type subunit 2A; −5.8 lfc, P=0.001) followed by GPR1 (G protein coupled receptor 1, high-binding affinity receptor for the adipokine Chemerin) (−5.8 lfc, P=0.0001) and TIDG4 (Tigger transposable element-derived protein 4) (−5.5 lfc, P=0.02). The most significantly downregulated gene was ATP2B2 (ATPase plasma membrane ca^2+^ transporting 2; 3.4 lfc, p<0.0001).

### Skeletal muscle transcriptome response to CE

A list of the top 20 upregulated genes is shown in Table 3, Of these 20, 5 also appear in the list of the top 20 upregulated genes following AE (CHI3L1, MT1A, SFN, C3orf52, and SOC3). The gene with the largest fold increase in response to CE was stratifin (SFN) (6.8 lfc, P=0.001), which has several roles relating to muscle function (13). Other genes with large fold increases were metallothionein 1A (5.8 lfc, P=0.01), which has roles in maintaining cellular health (14), and Chitinase-3 like protein-1 (CHI3L1, 5.6 lfc, P=0.002), a glycoprotein that mediates inflammation. The most significantly upregulated gene following CE was HMOX1 (heme oxygenase 1; 3.8 lfc, P<0.0001), which has been shown to be a vital gene in the adaptation to aerobic exercise and to maintain skeletal muscle health during endurance training (15). A list of the top 20 downregulated genes is shown in Table S2. The gene with the largest fold decrease was UNC13C (unc-13 homolog C; −2.4 lfc, P=0.01), followed by ABRA (Actin Binding Rho Activating Protein; −2.3 lfc, P0.009) and FLRT3 (fibronectin leucine rich transmembrane protein 3; −2.2, P=0.02). The most significantly downregulated gene was SDC4 (syndecan 4), a cell surface proteoglycan (−1.4 lfc, P=0.001) crucial for muscle differentiation (16).

**Table 3.** Top 20 upregulated genes following CE.

| Gene Symbol | Gene name | Gene Function | Log2fc | Adj P value |
| --- | --- | --- | --- | --- |
| SFN | Stratifin | Has a crucial in maintaining cell integrity. Functions include cell growth and development, cell cycle regulation, response to cellular stress, a scaffold for protein interactions. In muscle, it modulates oxidative stress, creating a favourable environment for muscle repair and regeneration. | 6.8 | 0.001 |
| MT1A | Metallothionein 1A | Has roles in protection against reactive oxygen species, regulating physiological balance and immune homeostasis | 5.8 | 0.01 |
| CHI3L1 | Chitinase 3 like 1 | A glycoprotein that plays a role in inflammatory and immunological disease, including cancer, Alzheimer's disease and atherosclerosis. It is upregulated in response to tissue injury or inflammation potentially influencing the repair and regeneration of muscle fibres by regulating the activity of immune cells like macrophages. | 5.6 | 0.002 |
| ADAMTS17 | ADAM metallopeptidase with thrombospondin type 1 motif 17 | A member of the ADAMTS family of enzymes, which are involved in the degradation of extracellular matrix components. They have important roles in tissue remodeling, growth, and repair, especially in processes like wound healing, development, and pathological conditions such as fibrosis and arthritis. | 4.8 | 0.01 |
| SOCS3 | Suppressor of cytokine signalling 3 | Negative regulator of cytokine signalling, inhibiting pro-inflammatory cytokines such as IL-6, IL-10 and IL-4. | 4.6 | 0.008 |
| LRP8 | LDL receptor related protein 8 | A member of the LDL receptor-related protein family with roles in lipid metabolism, cell signaling, and neuronal development. In muscle it has been implicated in the regulation of development and regeneration, potentially by modulating the interactions between muscle cells and the extracellular matrix. | 4.5 | 0.003 |
| C3orf52 | Chromosome 3 open reading frame | Precise biological functions are not yet fully elucidated | 4.3 | 0.01 |
| PTX3 | Pentraxin 3 | Plays a role in the innate immune system, inflammation and tissue repair | 4.1 | 0.01 |
| UBASH3B | Ubiquitin associated and SH3 domain containing B | Also known as STAP1 (Signal Transducing Adaptor Protein 1). It is, plays a key role in immune cell activation and response, with potential effects on muscle repair and regeneration. | 4.0 | 0.0002 |
| GADD45A | Growth arrest and DNA damage inducible alpha | Involved in cell cycle regulation, apoptosis, and DNA repair. In muscle it plays a role in repair, and regeneration with increases in expression seen during periods of muscle injury or inflammation. | 3.9 | 0.01 |
| HMOX1 | Heme oxygenase 1 | An enzyme that catalyzes the degradation of heme into biliverdin, carbon monoxide (CO), and iron. In muscle, it protects against oxidative stress. | 3.9 | 1.867E-13 |
| TNC | Tenascin | A family of extracellular matrix (ECM) proteins that play key roles in tissue architecture, cellular signaling, and the regulation of cell behaviour. It has a key role in muscle repair following injury. | 3.8 | 7.08E-07 |
| NNMT | Nicotinamide N-Methyltransferase | Plays an important role in NAD<sup>+</sup> metabolism by controlling the availability of nicotinamide. NAD<sup>+</sup> is a crucial coenzyme in the electron transport chain for ATP production in muscle cells. It also acts as a substrate for sirtuins, enzymes that regulate cellular stress response, aging, and mitochondrial function. | 3.8 | 0.04 |
| RUNX1 | Runt related transcription factor 1 | Plays a role in regulating the differentiation of satellite cells into myotubes and mature muscle fibres via regulation of myogenic transcription factors such as MyoD and myogenin. | 3.7 | 0.005 |
| SERPINB1 | Serpin family B member 1 | A serine protease inhibitor that plays a crucial role in modulating inflammatory responses and protecting tissues from excessive proteolytic damage | 3.7 | 0.005 |
| ACTC1 | Actin, alpha, cardiac muscle 1 | Also known as cardiac actin. It is a type of actin protein that plays a crucial role in the structure and function of muscle cells, particularly in cardiac muscle. It is a major component of the actin filaments that are part of the cytoskeleton and contribute to muscle contraction and cellular structure. | 3.6 | 4.90E-07 |
| ARMC9 | Armadillo repeat containing 9 | A ciliary protein, playing a role in the function of primary cilia, which are sensory organelles involved in receiving and processing signals. It has not been widely studied in skeletal muscle | 3.6 | 0.04 |
| TCAF2 | TRPM8 channel associated factor 2 | involved in various biochemical processes, particularly in folate metabolism and cell signaling. | 3.6 | 0.007 |
| TIMP1 | TIMP metalloproteinase inhibitor 1 | Plays a critical role in regulating the activity of matrix metalloproteinases, a group of enzymes responsible for the breakdown of extracellular matrix components. It plays a critical role in maintaining the structural integrity and mechanical properties of muscle tissue, and has roles in muscle repair and regeneration. | 3.5 | 3.77E-07 |
| SERPINE1 | Serpin family E member 1 | Also known as plasminogen activator inhibitor-1 (PAI-1), is a protein that plays a crucial role in regulating fibrinolysis. In muscle it plays a key role in tissue remodeling by controlling ECM degradation and matrix turnover | 3.5 | 0.001 |

Table represents the log fold change values from baseline to 24h following an unaccustomed session of combined resistance and aerobic exercise (CE) together with a brief description of known function. Abbreviations: fc, fold change

### Intramuscular inflammatory response to exercise

KEGG enrichment analysis showed that the pathways enriched 24h following bouts of either AE or CE were dominated by those involved in inflammation. These included the TNF-alpha signalling pathway, NF-kappa B pathway, MAPK signalling pathway, and the IL-17 signalling pathway (Figures 2A and 3A). These pathways play critical roles in orchestrating the immune response following skeletal muscle injury. Upregulated genes within these pathways following CE include CXCL1, CXCL2, CXCL3 and CCL2 that are known to recruit neutrophils and monocytes, suggest active leukocyte infiltration into the muscle. This is essential for clearing cellular debris and initiating tissue remodelling. Furthermore, increased expression of key receptors such as IL-1 and IL-17 receptors highlights the activation of signaling cascades that propagate the inflammatory response. IL1R1, a receptor for IL-1, plays a pivotal role in amplifying pro-inflammatory cytokine signaling, while IL17RE mediates IL-17-driven immune responses, which are crucial for defence and tissue repair. CHI3L1 (Chitinase-3-like 1), one of the most upregulated genes following both AE and CE, is a glycoprotein involved in macrophage polarization and protection against inflammation (17). A similar inflammatory profile was seen following AE with the addition of upregulation of the chemokine and toll-like receptor signalling pathways and TH17 cell differentiation processes. Within these pathways, DEG’s included additional chemokines (CXCL10, CXCL11, CCL8, CCL19, CXCR4), which play critical roles in guiding leukocyte migration and enhancing immune cell recruitment to sites of injury (18). Members of the signal transducer and activator of transcription (STAT) protein family, (STAT1, STAT3, STAT5A, STAT5B), JAK1, activator of the STAT proteins, and JAK3 highlighting their involvement in cytokine-driven signaling cascades that regulate inflammation and muscle regeneration via effects upon satellite cells (19). Together, these findings indicate a robust inflammatory signaling response to both AE and CE that orchestrates the recruitment and activation of immune cells while also laying the foundation for muscle regeneration. The KEGG and GO analyses underscore the dynamic interplay between pro-inflammatory chemokines and regulatory signaling proteins, suggesting a coordinated response aimed at resolving damage and supporting tissue remodeling in skeletal muscle.

**Figure 2.**
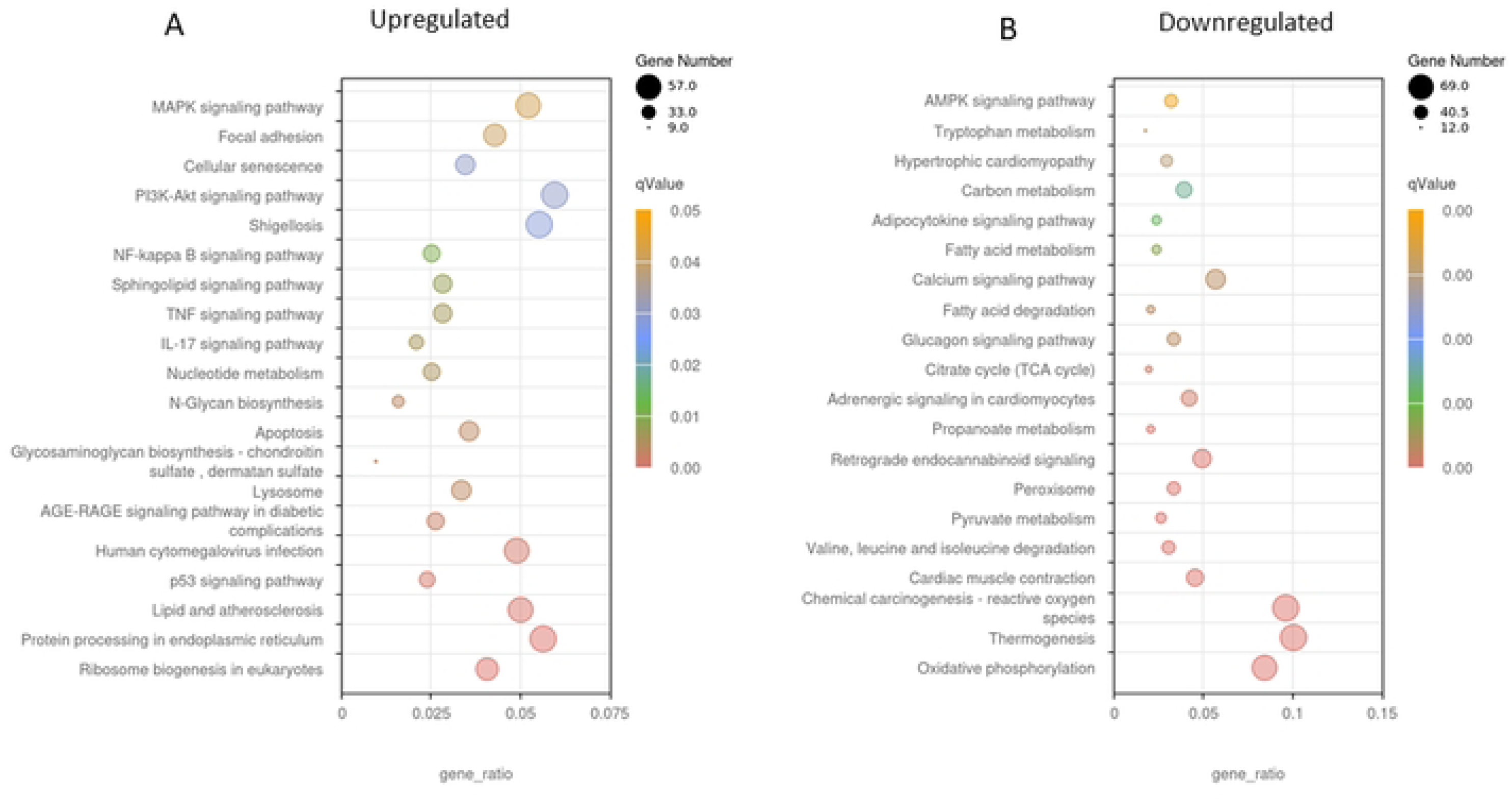
Kyoto Encyclopedia of Genes and Genomes (KEGG) analysis of DEG’s following unaccustomed CE. (A) processes upregulated 24h post CE and (B) processes downregulated 24h post CE. A high ‘proportion of enriched genes’ on the x-axis shows a high proportion of the pathway’s genes are enriched. The q value represents the false discovery rate and helps identify which GO terms or pathways remain significant after correcting for multiple comparisons. Darker colours (red) indicate a more highly significant result. The size of the dot relates to the proportion of enriched genes within the pathway. A smaller dot signifies fewer genes are enriched in the pathway, a large dot signifies many genes are enriched in this pathway.

**Figure 3.**
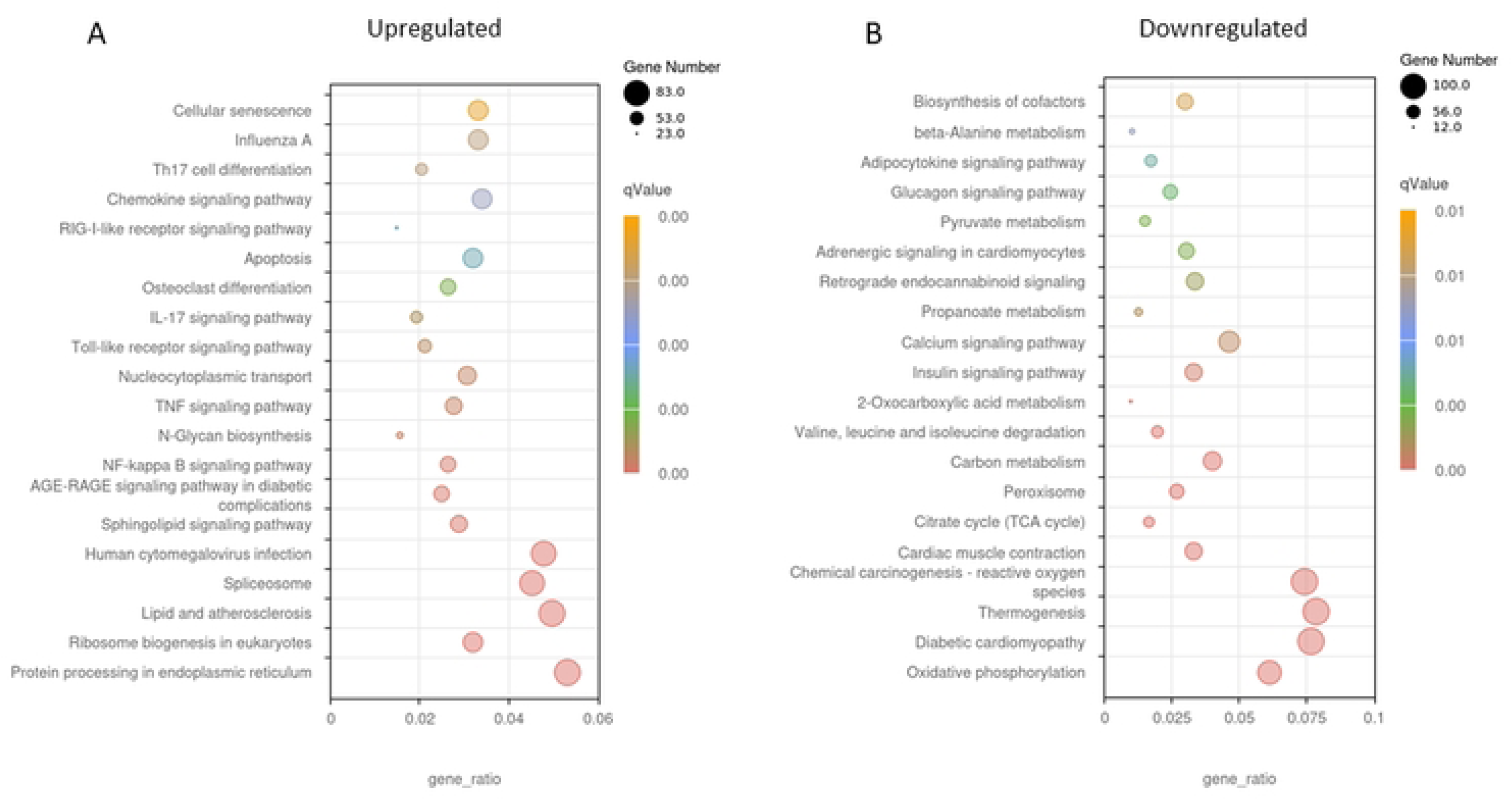
Kyoto Encyclopedia of Genes and Genomes (KEGG) analysis of DEG’s following unaccustomed AE. (A) processes upregulated 24h post AE and (B) processes downregulated 24h post AE. A high ‘proportion of enriched genes’ on the x-axis shows a high proportion of the pathway’s genes are enriched. The q value represents the false discovery rate and helps identify which GO terms or pathways remain significant after correcting for multiple comparisons. Darker colours (red) indicate a more highly significant result. The size of the dot relates to the proportion of enriched genes within the pathway. A smaller dot signifies fewer genes are enriched in the pathway, a large dot signifies many genes are enriched in this pathway.

### Regulation of mitochondrial biogenesis and translational machinery

In health, a clearly defined response to aerobic exercise is an increase in mitochondrial density via the initiation of mitochondrial biogenesis through PGC1-alpha. Here we report KEGG and GO analyses revealed significant downregulation of mitochondrial-related pathways following AE. In particular, master regulators of mitochondrial biogenesis were all significantly down-regulated 24 after AE such as PGC1-alpha and beta (PPARGC1A and B), and Perm1 (−1.9 lfc, P<0.0001; −2.1lfc, P<0.0001; −3.5lfc, P<0.0001respectively). This was seen together with a reduction in calcineurin expression (−1.1, P=3.65E-06). GO analysis revealed that multiple pathways relating to mitochondrial function/abundance and the respiratory chain were downregulated 24h following AE (Figure 5A); This was accompanied by reduced expression of genes encoding components of the respiratory chain, including NDUFA9 (complex I) and SDHA (complex II). A similar profile of genes was downregulated following CE (Figure 4B).

**Figure 4.**
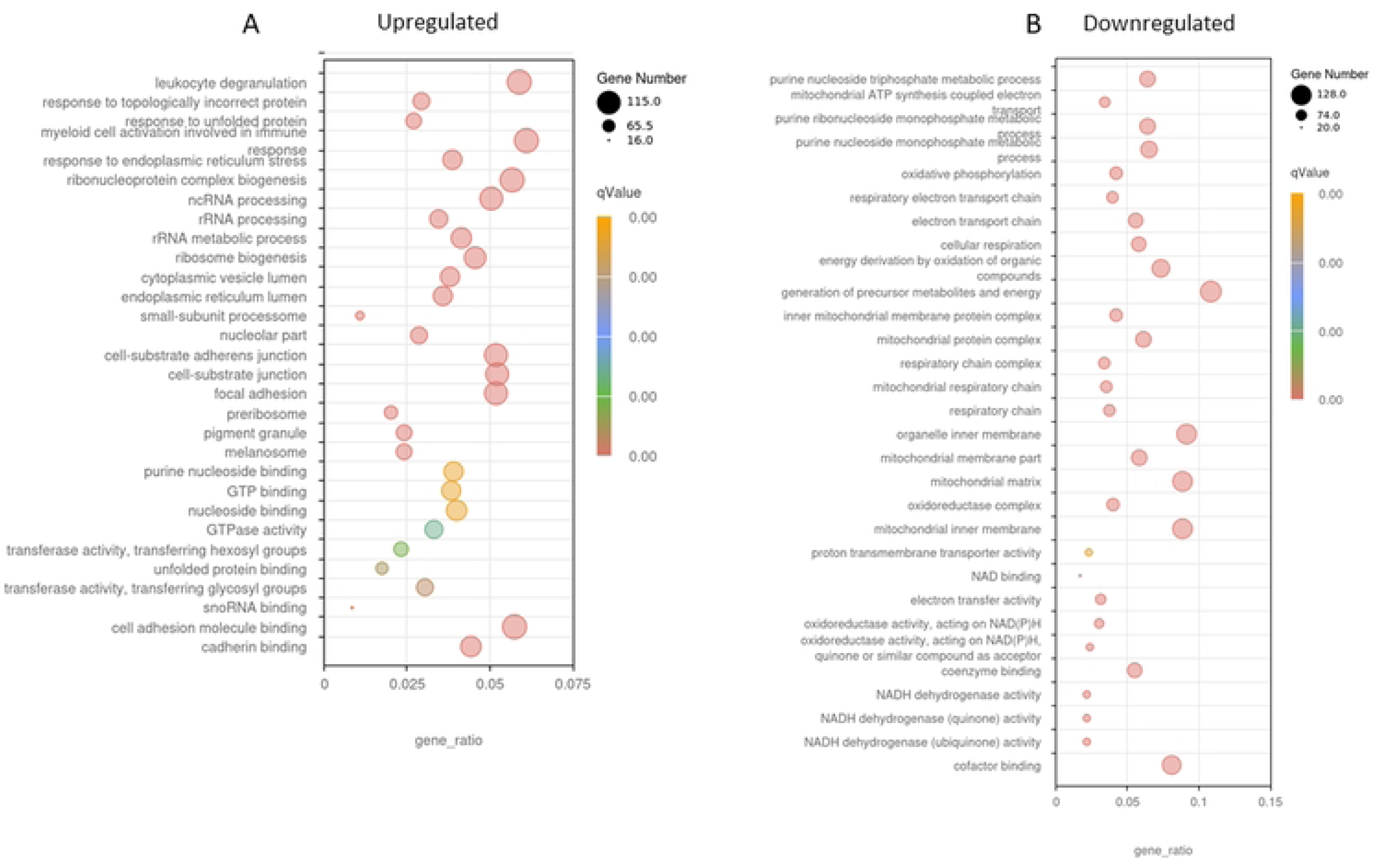
Gene Ontology (GO) analysis of DEG’s following unaccustomed CE. (A) processes upregulated 24h post CE and (B) processes downregulated 24h post CE. A high ‘proportion of enriched genes’ on the x-axis shows a high proportion of the pathway’s genes are enriched. The q value represents the false discovery rate and helps identify which GO terms or pathways remain significant after correcting for multiple comparisons. Darker colours (red) indicate a more highly significant result. The size of the dot relates to the proportion of enriched genes within the pathway. A smaller dot signifies fewer genes are enriched in the pathway, a large dot signifies many genes are enriched in this pathway.

**Figure 5.**
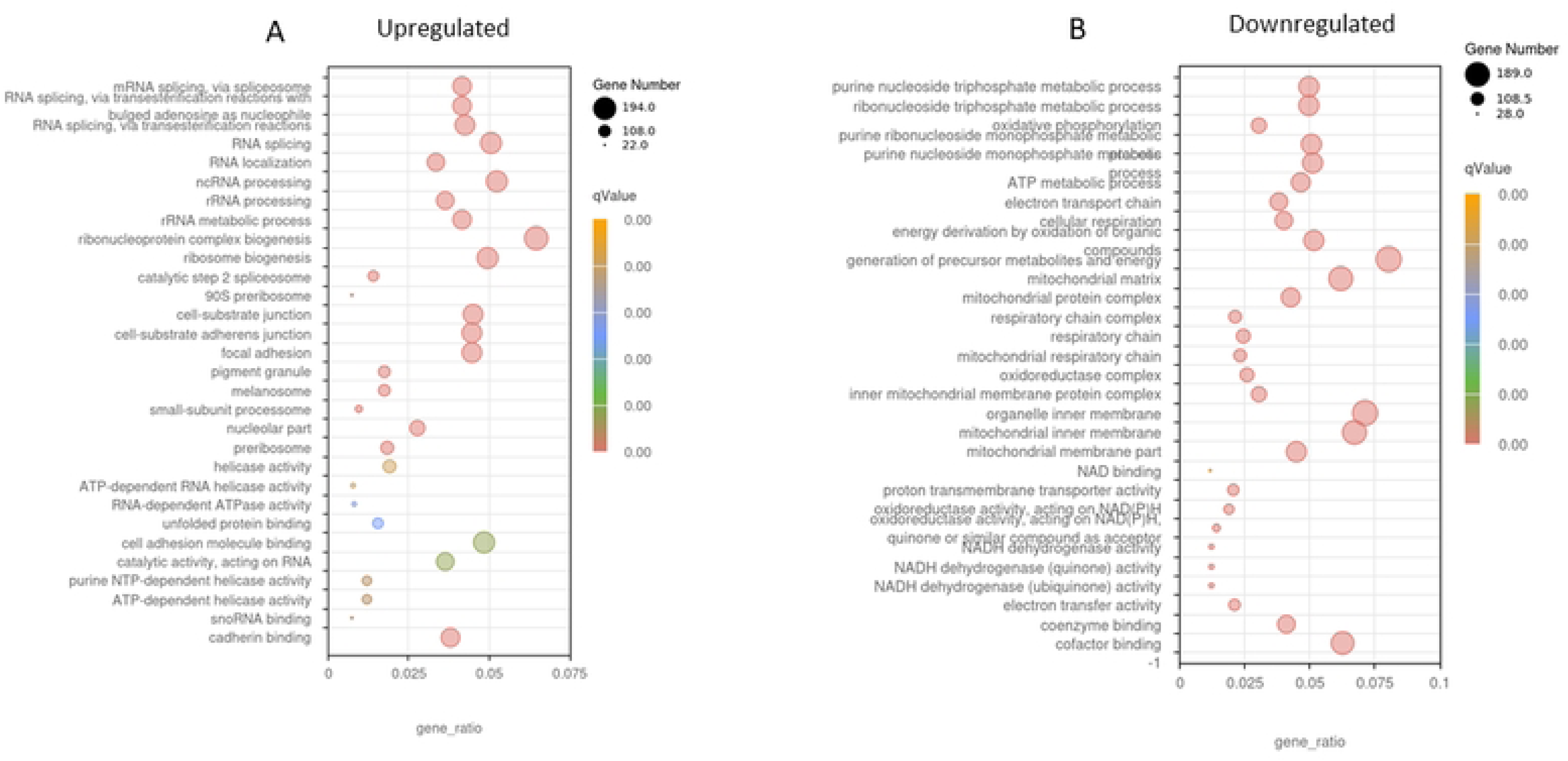
Gene Ontology (GO) analysis of DEG’s following unaccustomed AE. (A) processes upregulated 24h post AE and (B) processes downregulated 24h post AE. A high ‘proportion of enriched genes’ on the x-axis shows a high proportion of the pathway’s genes are enriched. The q value represents the false discovery rate and helps identify which GO terms or pathways remain significant after correcting for multiple comparisons. Darker colours (red) indicate a more highly significant result. The size of the dot relates to the proportion of enriched genes within the pathway. A smaller dot signifies fewer genes are enriched in the pathway, a large dot signifies many genes are enriched in this pathway.

### Anabolic/catabolic response to exercise

KEGG analysis (Figure2A) showed significant enrichment within the PI3k-Akt pathway following CE. This did not include any proteins involved in translation initiation, such as AKT1, RHEB, or mTORC1. Instead, there was a significant up regulation THBS1 (Thrombospondin 1; 3.5 lfc, P=0.0002) and OSMR (Oncostatin M Receptor; 2.4 lfc, P=0.002). Enrichment was also seen in the P53 signalling and the apoptosis pathways, which include catabolic related proteins CASP3 (caspase 3; 1.01 lfc, P=0.04), and GADD45A (growth arrest and DNA damage inducible α; 3.9lfc, P=0.01). There was also a significant increase in the expression of TRIM63, a muscle-specific E3 ligase. This has roles in the degradation and removal of damaged proteins following exercise. (1.5lfc, P=0.02). Following AE, enrichment was seen in the apoptosis pathway, but not in any of the other pathways seen following CE. In addition, AE resulted in a significant reduction in many genes within the insulin signalling pathway including IRS1, AKT2 and TSC2.

### Cellular senescence

KEGG analysis (Figures 2A and 3A) identified cellular senescence as one of the processes that was significantly upregulated following both AE and CE with a similar profile of genes upregulated in both groups. This included GADD45A, GADD45B, MYC, TP53, CDK2, NRAS, KRAS, MAPK11, CDK4, IL6, and CAPN2. In addition to these, AE promoted an increase in the expression of MRAS, NFKB1, CDKN1A, RASSF5, E2F4, TRAF3IP2, MCU, ENF3, RELA, RHEB, SLC25A6, ZFP36L2, CALM1, CALM2, PIK3R3, CCNA1 and TGFB3CE. CE also resulted in the upregulation of SERPINE1 which was not seen in the AE group.

## Discussion

In this study we performed full untargeted RNA sequencing of skeletal muscle biopsies from people with CKD before and 24h after an acute unaccustomed bout of either AE or CE in order to explore the exercise-induced changes in the skeletal muscle transcriptome. We found a large intramuscular inflammatory response to both forms of exercise and a downregulation in several processes relating to mitochondrial mass and function following AE. Understanding this acute response is vital to identify how the individual or population adapts to regular exercise. This will help us to design strategies to ensure optimal adaptation to exercise.

### Activation of intramuscular inflammation following exercise

Following both CE and AE there was a strong upregulation of pathways relating to inflammation. Intramuscular inflammation is an expected response to unaccustomed exercise that can create a degree of muscle damage (20). This process serves to remove damaged proteins within the injury site and is required for proper recovery and repair. Here we report an increase in the gene expression of the neutrophil attractant chemokines CXCL1, CXCL2, CXCL3 and CXCL4 after both forms of exercise. In the early stage of the injury response, neutrophils are attracted to the injury site where they act to clear cellular debris by phagocytosis and cell lysis, and further propagate the inflammatory response by cytokine secretion. We also saw increases in the expression of CCL2, a potent chemotactic factor for monocytes, which results in an increased abundance of macrophages at the injury site to aid repair. This data suggests an infiltration of leukocytes into the muscle occurred following the exercise. This is a well-characterised response, key in the processes of adaptation to exercise, and is one that we have previously described in the CKD population (6, 7). Interestingly, this response was seen after both modes of exercise suggesting that in this population AE is sufficient to confer a small amount of muscle damage eliciting an inflammatory response. One of the most highly upregulated genes following both AE and CE was CHI3L1, a glycoprotein that mediates inflammation and macrophage polarisation. It has been shown to protect skeletal muscle from TNF-alpha induced inflammation (21), and therefore may be upregulated as a protective response to the large inflammatory reaction to the exercise. AE promoted an increase in the expression of IL-6, a well-characterised response to skeletal muscle contraction. Exercise-induced IL-6 helps to initiate the anti-inflammatory effects of exercise, fuel mobilisation during exercise and reduces adiposity (22). This is also an effect that we have described previously in this population (6).

Following AE, we also saw an upregulation in the expression of several genes within the STAT family, which are activated by a wide range of cytokines, including IL-6. STAT3 is known to be phosphorylated following resistance exercise (23), resulting in its translocation to the nucleus where it regulates target gene expression. It has been shown to have a role in muscle regeneration via effects upon satellite cell proliferation and differentiation (24). STAT5A was upregulated following both modes of exercise, and whilst there is not as much data on the role of STAT5A, it has been shown to be upregulated following aerobic exercise before (25). *In vitro* studies have shown that it promotes IGF-1 expression in C2C12 cells (26) and therefore it may have a role to play in hypertrophy.

What we are unable to conclude from this data is for how long this inflammatory response remains. If this response in CKD patients is exaggerated or prolonged, it may impede muscle recovery and adaptation.

### Mitochondrial responses to exercise

It is well-characterised that several forms of exercise result in skeletal muscle mitochondrial biogenesis, the synthesis of new mitochondrial organelles, which has also been demonstrated in the CKD population via analysis of mtDNA copy number (27) via analysis of mtDNA copy number. However, in this study we report a down-regulation in processes relating to mitochondrial function/abundance and the respiratory transport chain 24 hours after a session of either aerobic exercise alone or combined exercise. This was seen alongside downregulation in the gene expression of PGC1-alpha and beta (PPARGC1A and B) and Perm1, master regulators of mitochondrial biogenesis suggesting that many aspects of mitochondrial function/health, 24h following both forms of exercise. These findings align with our previous observations (8), which highlighted an abnormal mitochondrial response to exercise training in this population. In that earlier study, which also used muscle biopsies from the ExTra CKD study, we found no effect of exercise training on mitochondrial abundance in individuals with CKD. If this absence of a biogenesis response continues, it could explain why we reported no increase in mitochondrial protein levels after training (8). Analysis of biopsies taken after exercise following a period of training would be required to answer this question. It is important to discuss this observation in the context of the biopsy timing. The activation of PCG-1alpha following exercise may be reasonably short-lived, with expression returning to baseline within 10h of exercise (28), although it has been shown to have remained significantly elevated up to 24h post-exercise (29). There was also a strong downregulation of many of the structural components of the mitochondria which is in line with this reduction in mitochondrial biogenesis. To fully understand the implications of these observations, a time course analysis of the effect of exercise on the upregulation of mitochondrial-related processes should be performed, including much earlier time points post-exercise before we can reliably conclude a dysregulated mitochondrial response to exercise in this population. Mitochondrial dysfunction and inflammation are deeply interconnected processes, with the inflammatory response reported here potentially contributing to and exacerbating mitochondrial dysfunction. This mitochondrial dysfunction may partly explain the poor exercise tolerance and low exercise capacity seen in these patients.

### Effects on skeletal muscle protein synthesis and degradation pathways

KEGG and GO analyses showed significant enrichment within the PI3k-Akt pathway following CE. However, this did not include any of the proteins involved in the canonical pathway of protein synthesis and instead included proteins that have previously been shown to have catabolic roles within skeletal muscle. For example, THBS1 was strongly upregulated which has been shown to play a role in skeletal muscle atrophy via effects on TGF-b-Smad-2 signalling activating both the autophagy-lysosomal and the ubiquitin-proteasome system (30). Increased expression of THBS1 has been reported following overload previously (31), which the authors concluded was involved in exercise-induced angiogenesis, another known role for this protein. Angiogenesis is not a process that was seen to be significantly enriched, therefore the true effect of upregulation of this gene is unknown. Onconstatin M (OSMR) was also seen to be significantly upregulated following CE. OSMR is a cytokine of the interleukin-6 family and is a potent inducer of muscle atrophy (32) via the JAK/STAT3 pathway. Mice lacking OSMR are protected from atrophy (32). In a mouse model, prolonged expression of OSMR had an inhibitory effect on skeletal muscle regeneration following injury again via the JAK1/STAT1/STAT3 pathway (33). This suggests that in the CKD population there may be an impaired response to muscle injury via OSMR, but this needs further investigation. TRIM63, a muscle-specific E3 ligase, was upregulated, an effect that we have previously shown in this population (7) as was CASP3, caspase 3, which is involved in the initial step of actin and myosin degradation. These genes are likely to have key roles in the removal of damaged proteins following exercise and therefore in the processes of repair and adaptation. Upregulation of these genes may be in response to the large inflammatory reaction; The NF-kB pathway is activated by pro-inflammatory cytokines such as TNF-alpha and in turn can upregulate TRIM63 and consequently protein degradation. In summary, 24h post CE the transcriptome response largely involves atrophy-related processes rather than anabolic. This observation may simply represent a high turnover of skeletal muscle protein following damage leading to muscle repair. This is an essential part of the remodelling process following exercise serving to clear damaged proteins replacing them with functional ones which supports adaptation, and is likely to be highly inter-related to the inflammatory response seen. As discussed above, this picture may simply reflect the time point of the biopsy and needs further validation.

Following AE there was a significant downregulation of many genes that encode for proteins within the PI3k-Akt pathway including IRS1, AKT2 and TSC2. However, this was not seen following CE. Without biopsies at earlier time points, it is impossible to know if there was an anabolic response that was missed, or if CKD perturbs this expected anabolic response in some way. Finally, we also reported an increase in growth arrest and DNA damage-inducible alpha (GADD45A), which has been shown to reduce mitochondrial abundance and oxidative capacity resulting in atrophy of type II fibres in mice, which led to a reduction in strength, force production and exercise capacity (34). The mRNA expression of GADD45A is positively associated with muscle weakness in humans (34), and its expression increases during prolonged periods of physical inactivity (35). It has been described as an atrophy inducer factor and highlighted as a potential target to treat muscle weakness in humans (34). Interrogation of the MetaMEx database (22) shows that upregulation of GADD54A following both aerobic and resistance exercise has been shown several times previously, but all these studies took muscle biopsies much earlier post-exercise (1-4h), with no studies reporting as far out as 24h. Therefore, its biological function in this context is unknown. However, given its integral role in skeletal muscle atrophy, this warrants closer investigation.

### Cellular Senescence

Cellular senescence was identified by KEGG analysis as being strongly up-regulated by both AE and CE. This is a condition of permeant cellular proliferation arrest and is upregulated by several different environmental conditions. CE upregulated Serpine-1 (also known as plasminogen activator inhibitor - 1 (PAI-1)), which KEGG pathway analysis shows has a role in paracrine senescence. It can be stimulated by several signalling cascades including pro-inflammatory and pro-fibrotic, and also plays a key role in extracellular matrix remodelling and the development of fibrosis. PAI-1 inhibitors attenuate atrophy and increase strength in animal models of sarcopenia (36). GADD45A as described above, also plays a role in cellular senescence through interactions with Cyclin-dependent kinase 1 (CDK1) and p21 (aka. CDKN1A), which directly inhibits the cell cycle machinery (37). Cyclin-dependent kinase 2 (CDK2) was also upregulated following both AE and CE. CKD2 has been shown to phosphorylate the myogenic factor MyoD preventing skeletal muscle differentiation (38), a key process in skeletal muscle repair and regeneration and in promoting satellite cells out of senescence. We did not observe any increases in the expression of any of the key genes involved in skeletal muscle myogenesis such as MyoD, myogenin or Myf5, which suggests that this process was not activated following either exercise type at the time points we studied and again, may suggest that CKD patients have an impaired muscle regeneration response following exercise. This requires much more investigation and may also be linked to the intramuscular inflammatory response (39). Acute exercise has been previously shown to promote senescence in fibro-adipogenic progenitor cells (40), and this has been shown to be required for effective skeletal muscle regeneration. Bulk RNA sequencing does not allow us to identify in which cell types this process was upregulated, which highlights the importance of single-cell RNA sequencing and spatial approaches to tease apart the effects in different cell types to further our understanding of the tissue-specific response to a bout of exercise in those with CKD. Research has shown that regular exercise training reduces the number of senescent cells by activating those immune cells that are responsible for their clearance (41). It would be interesting to know if this reduction in the senescent cell population within skeletal muscle is also seen following exercise training in the CKD population.

### Other observations

Interestingly the number of DEG’s was very different between groups, with a higher number of DEG’s seen following AE compared to CE. This is the opposite to published data (42), where a higher number of responsive genes were found following resistance exercise compared to aerobic exercise. We suggest this may reflect the overload stimulus used at this very early stage of the exercise programme in the CE arm in these deconditioned patients may have been insufficient to induce a large change in the transcriptome.

Some other interesting and previously unreported observations in this population were made. The most upregulated gene following CE was stratifin, found in stratified eukaryote cells. This protein has been associated with wound healing in skeletal muscle (13) and is therefore likely to be part of the remodelling response to the exercise. It is also a potent activator of Nrf2 and plays an important role in preventing oxidative stress by the activation of the Nrf2 signalling pathway. It has been shown to protect skeletal muscle against damage induced by exhaustive exercise (43). HMOX1 was seen to be upregulated following both AE and CE. This has been shown to be a vital gene in the adaptation to aerobic exercise and to maintain skeletal muscle health during endurance training (15). This enzyme is activated by an increased level of heme within skeletal muscle due to microtrauma created by the exercise. It has a role in reducing oxidative stress and upregulation of HMOX1 can protect against injury (15). It enables skeletal muscle adaptation via effects on satellite cell activation, fibre type transition and mitochondrial function (15). Overall, it has been described as a central regulator in the physiologic response of skeletal muscle to exercise and has been indicated to be a target for skeletal muscle atrophy (44).

There was a significant amount of overlap in the response to AE and CE, with 25% of the top 20 upregulated genes seen following both AE and CE. It has been suggested that an acute unaccustomed bout of exercise irrespective of the mode of exercise may largely reflect a general stress response within skeletal muscle rather than a targeted response to a specific form of exercise (45), which might explain the large degree of similarity between the two forms of exercise.

### Limitations

As already discussed throughout, these results must be interpreted within the time frame that the biopsy was taken, which represents just a single time point of a very dynamic process. Many changes in gene expression are very transient (return to baseline levels within hours after the exercise bout), so using the 24h timepoint following exercise it is likely that we missed some key acute changes and what we present here represents the sustained or delayed response to the exercise. To fully understand the molecular response to exercise in these patients, future studies should take serial biopsies starting at much earlier time points post-exercise. This is a difficult study to do in clinical populations and therefore an alternative might be to use a primary cell culture model of human CKD skeletal model and mechanically stretch these cells *in vitro* collecting the cells at various time points post stretch to understand the full-time course of the skeletal muscle transcriptome response to exercise, which of course has its own inherent limitations. Another important limitation is that we did not include a healthy control group in this analysis, so we are unable to infer what is a CKD-specific response to the exercise, and what simply represents a normal response. However, a lot of data is available on the skeletal muscle transcriptomic response to exercise in various populations, which has been used for comparison in the discussion here. It is important to point out that there is a difference between the groups for BMI. The AE group had a BMI of 27.5 (overweight) and the CE group had a BMI of 32.5 (obese). Data suggests that obesity can alter the skeletal muscle molecular response to exercise (46), notably glycogen synthase kinase-3β signaling. Therefore, it is hard to determine if the differences in responses seen here are due to differences in BMI, or the different modes of exercise. Given the high rates of obesity in people with CKD, this is an important avenue for future research. This data is based upon a small sample size of 4 participants per group which likely limits statistical power and the generalisability of the findings. It would be important to repeat this in a larger group to confirm the responses we have seen here. Finally, these results require further validation. We were unfortunately unable to perform experimental verification due to lack of sample availability. Moving forwards, in order to understand the functional relevance of these changes in gene expression and the role they play in the exercise response, functional validation is required. It is also important to note that we have not performed an integrated proteomic analysis, therefore the true biological meaning of these transcriptional changes in unknown.

## Conclusion

For the first time, we report here the change to the skeletal muscle transcriptome 24h after an unaccustomed bout of aerobic or combined exercise in the CKD population. The main observation was a large intramuscular inflammatory response to both forms of exercise, with what may be an impaired muscle regenerative response. A downregulation in several processes relating to mitochondrial mass and function following AE was surprising, but supports our previous data in this population (8) and possibly highlights the mitochondria as a therapeutic target for improving adaptation to exercise with agents such as resveratrol to increase mitochondrial biogenesis. However, due to the timings of the biopsy collections, it is not known what the early response to exercise was in many of these processes, and this is an important limitation of the current data. A study including earlier biopsy time points is crucial for the complete understanding of the response of the skeletal muscle transcriptome to exercise in patients with CKD. Furthermore, this study only investigated responses within CKD stages 3b-5. It would be of great interest to compare these responses with those individuals at earlier stages of CKD, and those receiving renal replacement therapy.

## Acknowledgements

This is a summary of independent research funded by the Leicester Hospitals Charity Kidney Care Appeal and the Stoneygate Trust, and supported by the National Institute for Health and Care Research (NIHR) Leicester Biomedical Research Centre (BRC). The views expressed are those of the author(s) and not necessarily those of the Kidney Care Appeal, the Stoneygate Trust, the NIHR or the Department of Health and Social Care. We thank Dr Douglas Gould and Dr Soteris Xenophontos for their help with the collection of the muscle biopsy samples. For the purpose of open access, the author has applied a Creative Commons Attribution license (CC BY) to any Author Accepted Manuscript version arising from this submission. There are no conflicts of interest to disclose.

